# Fermentative *Escherichia coli* makes a substantial contribution to H_2_ production in coculture with phototrophic *Rhodopseudomonas palustris*

**DOI:** 10.1101/610568

**Authors:** Amee A. Sangani, Alexandra L. McCully, Breah LaSarre, James B. McKinlay

**Affiliations:** Department of Biology, Indiana University, Bloomington, IN; Biology Undergraduate Program, Indiana University, Bloomington, IN; Department of Civil and Environmental Engineering, Stanford University, CA

**Keywords:** Coculture, Hydrogen, Biofuel, Cross-feeding, Syntrophy, Rhodopseudomonas palustris, Formate hydrogenlyase

## Abstract

Individual species within microbial communities can combine their attributes to produce services that benefit society, such as the transformation of renewable resources into valuable chemicals. Under defined genetic and environmental conditions, fermentative *Escherichia coli* and phototrophic *Rhodopseudomonas palustris* exchange essential carbon and nitrogen, respectively, to establish a mutualistic relationship. In this relationship, each species produces H_2_ biofuel as a byproduct of their metabolism. However, the extent to which each species contributes to H_2_ production and the factors that influence their relative contributions were previously unknown. By comparing H_2_ yields in cocultures pairing *R. palustris* with either wild-type *E. coli* or a formate hydrogenlyase mutant that is incapable of H_2_ production, we determined the relative contribution of each species to total H_2_ production. Our results indicate that *E. coli* contributes between 32% and 86% of the H_2_ produced in coculture depending on the level of ammonium excreted by the *R. palustris* partner. An *R. palustris* strain that stimulated rapid *E. coli* growth through a high level of ammonium excretion resulted in earlier accumulation of formate and acidic conditions that allowed *E. coli* to be the major contributor to H_2_ production.

## Introduction

The collective activities of microbial communities can be harnessed to benefit society in ways ranging from the degradation of pollutants to the production of valuable chemicals and fuels (Zuroff & Curtis, 2012, Johns *et al.*, 2016, Cavaliere *et al.*, 2017). Synthetic communities (i.e., cocultures) pairing fermentative and phototrophic purple nonsulfur bacteria have long been viewed as an attractive means by which to convert carbohydrates to hydrogen gas (H_2_) biofuel (Odom & Wall, 1983). In such communities, fermentative bacteria often dedicate some electrons extracted from carbohydrates to H_2_ production, but most of the electrons end up in excreted organic acids and alcohols. Purple nonsulfur bacteria can use energy from light to access electrons in fermentation products and use these electrons for both biosynthesis and H_2_ production, thereby increasing the total H_2_ yield. While the advantages of such communities for achieving a higher H_2_ yield in a consolidated process have long been known (Odom & Wall, 1983), the contributions of each species to H_2_ production and the underlying microbial interactions have been difficult to characterize due to a lack of reproducibility and tractability in these communities.

We previously developed a highly reproducible and tractable coculture between fermentative *Escherichia coli* and the purple nonsulfur bacterium, *Rhodopseudomonas palustris* (LaSarre *et al.*, 2017). As in previous cocultures of this kind, *E. coli* ferments glucose into excreted organic acids and ethanol. One organic acid, formate, can be further converted by *E. coli* into H_2_ and CO2 via formate hydrogenlyase (FHL) (Pinske & Sawers, 2016). Formate cannot be metabolized by *R. palustris,* but *R. palustris* readily consumes the other organic acids, namely acetate, lactate, and succinate. *R. palustris* does not metabolize sugars and is thus reliant on *E. coli* for the carbon and electrons in these organic acids (LaSarre *et al.*, 2017). Stable coexistence of the species in coculture is assured by requiring that *E. coli* rely on *R. palustris* for essential nitrogen. This dependency is achieved by (i) providing N_2_ gas as the sole nitrogen source, as only *R. palustris* can convert N_2_ into NH_4_^+^ via nitrogenase, and (ii) by using *R. palustris* mutants that excrete NH_4_^+^ as a nitrogen source for *E. coli* (LaSarre *et al.*, 2017). These conditions also drive H_2_ production by *R. palustris,* as H_2_ an obligate byproduct of the nitrogenase reaction.

Here we use an *E. coli* mutant lacking FHL activity to assess the contribution of each species to H_2_ production in coculture. We find that either species can make the majority contribution to H_2_ production depending on the level of NH_4_^+^ excreted by the *R. palustris* partner. A highly cooperative *R. palustris* partner, exhibiting a high level of NH_4_^+^ excretion, leads to conditions that result in *E. coli* being the major contributor to H_2_ production.

## Materials and Methods

### Strains and growth conditions

All *R. palustris* strains were derived from the type strain CGA009 (Larimer *et al.*, 2004). *R. palustris* Nx (CGA4005) has a *nifA* mutation that results in NH_4_^+^ excretion during N_2_ fixation, a *hupS* deletion to prevent H_2_ oxidation, and a *uppE* deletion that prevents biofilm formation (Fritts *et al.*, 2017, LaSarre *et al.*, 2017). *R. palustris* NxΔAmtB (CGA4021) has the same mutations as the Nx strain and additional *amtB1* and *amtB2* deletions that result in 3-fold more NH_4_^+^ excretion than the Nx strain (LaSarre *et al.*, 2017). *E. coli* MG1655 was the wild-type (WT) strain (Blattner *et al.*, 1997). The ΔFdhF strain was created by transferring the *ΔfdhF: Km^R^* mutation from *E. coli* JW4040 (Baba *et al.*, 2006) into *E. coli* MG1655 using P1 phage transduction (Thomason *et al.*, 2007). Mutants were selected on LB agar with 30 μg/mL kanamycin and were verified by PCR. Empty vector (pCA24N) and the complementation vector (pCA24N_fdhF) (Kitagawa *et al.*, 2005) were electroporated into *E. coli* ΔFdhF and transformants were selected on LB agar with 25 μg/mL chloramphenicol. Monocultures and cocultures were grown in 10 mL of M9-derived coculture (MDC) medium (LaSarre *et al.*, 2017) in 27-mL anaerobic test tubes. Where appropriate, MOPS was added during the preparation of MDC medium using a 1M stock at pH 7 for a final concentration of 100 mM. Tubes were made anaerobic by bubbling with N_2_ and were then sealed with rubber stoppers and aluminum crimps, creating a headspace of 100% N_2_. MDC medium was supplemented with 1% v/v cation solution (100 mM MgSO4, 10 mM CaCl_2_) and 25 mM glucose. For *E. coli* monocultures, 15 mM NH_4_Cl was added. All anaerobic cultures were grown at 30°C, lying flat with shaking at 150 rpm beneath a 60 W incandescent bulb. Starter monocultures and cocultures were inoculated from single colonies suspended in 0.2 ml of MDC. Once fully grown, 0.1 mL of starter culture was inoculated into test conditions.

### Analytical Procedures

Cell densities were quantified by optical density at 660nm (OD660) using a Genesys 20 spectrophotometer (Thermo-Fisher). Growth curves used cell densities measured in the culture tubes. Growth rates were determined using values between 0.1 and 1.0 OD660 where there is linear correlation between OD660 and cell density. Growth yields were determined using OD660 values from initial and final time points measured in cuvettes, with samples diluted into the linear range as necessary. Glucose and soluble fermentation products were quantified using a Shimadzu high-performance liquid chromatograph as described (McKinlay *et al.*, 2005). H_2_ was quantified using a Shimadzu gas chromatograph as described (Huang *et al.*, 2010). To determine final pH values, whole cultures were centrifuged, the supernatants passed through 0.2 μm syringe filters, and the pH of the filtrate measured using a pH meter.

## Results

### Formate dehydrogenase-H is required for H_2_ production by *E. coli*

In cocultures pairing WT *E. coli* with *R. palustris* Nx, a mutant that excretes NH_4_^+^ during N_2_ fixation, both species are presumed to produce H_2_ (Fig 1). However, the contribution of each species to H_2_ production was unknown. To determine the contribution of each species to H_2_ production, we genetically disrupted H_2_ production in *E. coli*. We did not attempt to disrupt H_2_ production in *R. palustris* because H_2_ is an obligate byproduct of nitrogenase during the conversion of N_2_ to NH_4_^+^; consequently, *R. palustris* H_2_ production cannot be eliminated without simultaneously disrupting the NH_4_^+^ cross-feeding that underpins the mutualism. In *E. coli,* H_2_ is produced by the FHL complex, composed of formate dehydrogenase-H and hydrogenase-3, which converts formate to H_2_ and CO2 (Pinske & Sawers, 2016). Unlike other *E. coli* fermentation products, formate is not consumed by *R. palustris* (Fig. 1).

**Fig. 1.**
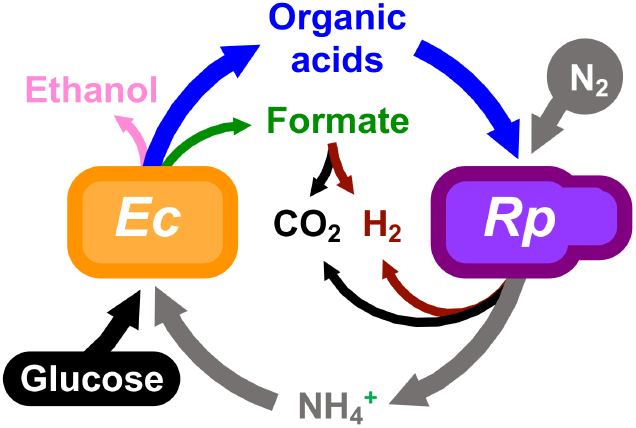
A mutualistic H_2_-producing coculture between WT *E. coli (Ec)* and *R. palustris (Rp)* Nx. *E. coli* ferments glucose into organic acids that serve as an essential carbon source for *R. palustris.* Ethanol and formate accumulate, but WT *E. coli* can convert formate into H_2_ and CO2 using FHL. *R. palustris* Nx converts N_2_ gas into NH_4_^+^ via nitrogenase and excretes some NH_4_^+^ that serves as an essential nitrogen source for *E. coli. R. palustris* produces H_2_ as a byproduct of the nitrogenase reaction.

It is well-known that *fdhF*, encoding formate dehydrogenase-H, is required for H_2_ production by *E. coli* (Pinske & Sawers, 2016). We therefore deleted *fdhF,* resulting in strain ΔFdhF, and examined the effects of the *fdhF* deletion on fermentative growth and metabolic trends in monoculture. The growth curves of WT and ΔFdhF *E. coli* monocultures were comparable (Fig. 2A), as were the growth yields (Fig. 2B) and growth rates (Fig. 2C). As expected, the ΔFdhF mutant did not produce any detectable H_2_ and had a higher formate yield than did the WT strain (Fig. 2D). Other fermentation product yields were comparable in the WT and ΔFdhF cultures (Fig. 2D). We verified that the loss of H_2_ production and higher formate yield in the ΔFdhF cultures were due to the lack of the *fdhF* gene and not due to polar effects of the deletion by complementing the ΔFdhF mutant with a plasmid bearing *fdhF* under an IPTG-inducible promoter (pCA24N_fdhF). Whereas the empty vector had no effect on product yields, complementation with pCA24N_fdhF restored H_2_ production and resulted in formate yields similar to WT levels (Fig. 2D). Complementation with pCA24N_fdhF also resulted in a higher lactate yield than the WT strain (Fig. 2D), though the reasons for this trend are not obvious. Because formate accumulation can acidify the culture medium we also measured the culture pH once growth had ceased. Despite the higher formate level in ΔFdhF cultures, there was no significant difference in final culture pH compared to WT cultures (Fig. 2E). Overall, all results verified that deletion of *fdhF* abolishes H_2_ production with a concomitant increase in formate yield but without affecting other growth and metabolic trends.

**Fig. 2.**
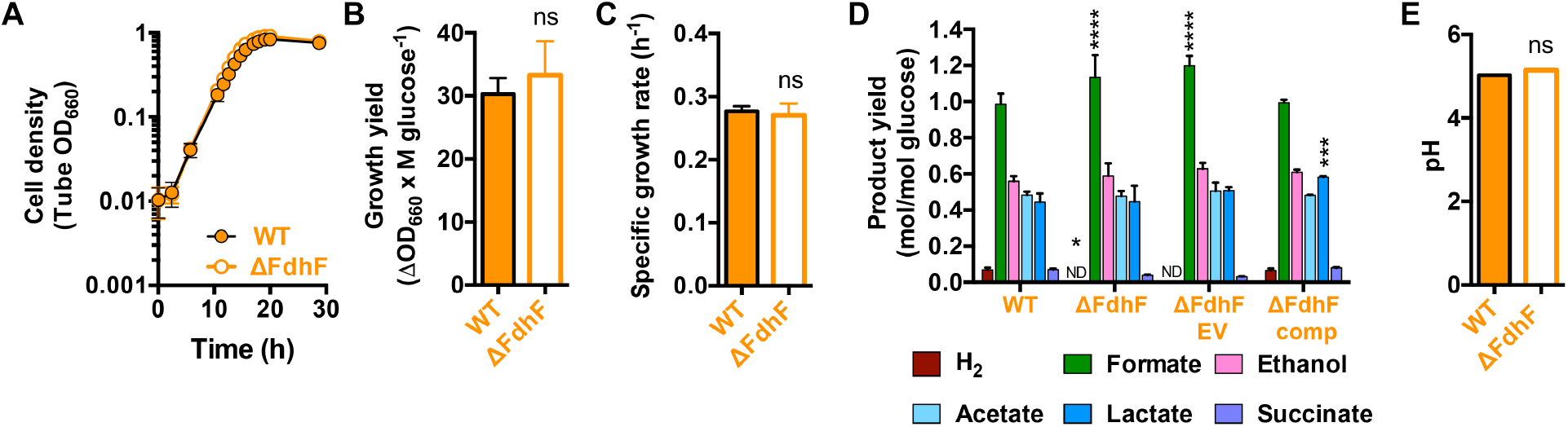
Growth and metabolic trends of WT *E. coli* and ΔFdhF mutant monocultures. Growth curves (n=3) (A), growth yields (n=6) (B), and growth rates (n=3) (C) of WT and ΔFdhF *E. coli* strains. (D) Fermentation product yields(n=6). Asterisks indicate a statistical difference from corresponding WT value with P<0.05 (*), P<0.001 (***), P<0.0001 (****), determined by two-way ANOVA with Sidak posttest. EV, empty vector (pCA24N); comp, complementation vector (pCA24N_fdhF). (E) Final pH (n=3). (A-E) Error bars, SD. (B, C, E) Statistical differences from WT trends were determined using an unpaired, two-tailed t test; ns, nonsignificant.

### *E. coli* contributes to H_2_ production in coculture with *R. palustris*

As deletion of *fdhF* abolished the conversion of formate to H_2_ by *E. coli* without altering other growth or metabolic trends, we deemed the ΔFdhF mutant suitable for assessing the contribution of *E. coli* to H_2_ production in coculture. We grew the ΔFdhF mutant with *R. palustris* Nx (i.e., Nx+ΔFdhF coculture) and compared growth and metabolic trends to cocultures pairing *R. palustris* Nx with WT *E. coli* (i.e., Nx+WT coculture). We observed no significant differences in the growth trends between the Nx+WT and Nx+ΔFdhF cocultures (Fig. 3A-C). The H_2_ yield in Nx+ΔFdhF cocultures was significantly lower than that of Nx+WT cocultures, whereas the formate yield was significantly higher (Fig. 3D). Despite that formate yields were significantly higher in Nx+ΔFdhF cocultures, the final pH was still similar to that of Nx+WT cocultures (Fig. 3E).

**Fig. 3.**
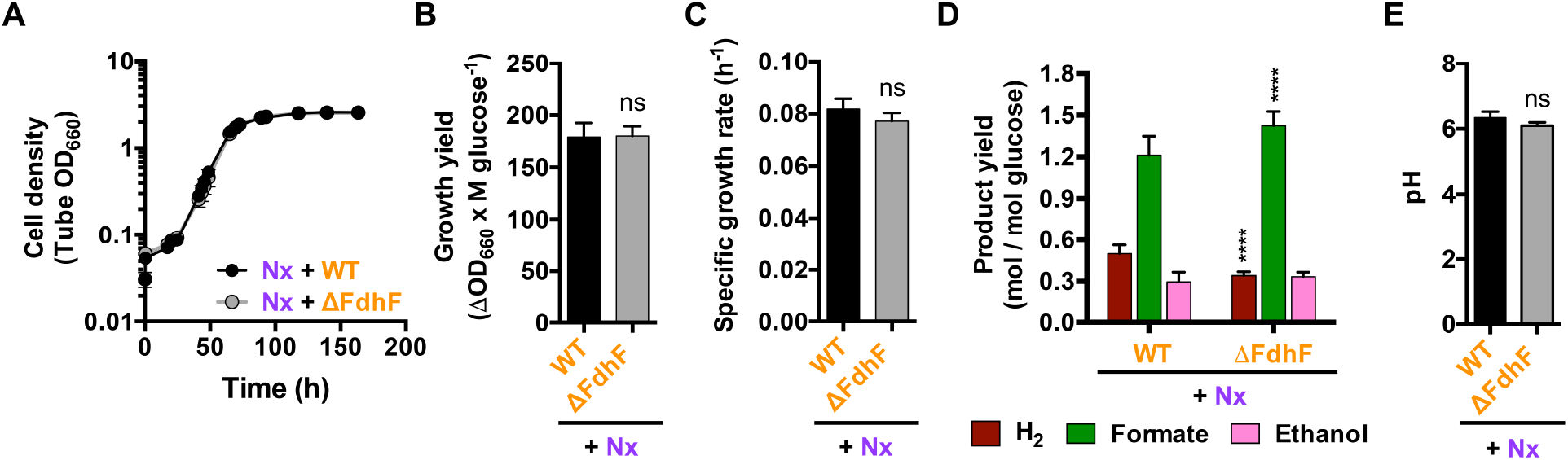
Growth and metabolic trends of cocultures pairing *R. palustris* Nx with either WT *E. coli* or the ΔFdhF mutant. Coculture growth curves (A), growth yields (B), growth rates (C), product yields (D), and final pH (E) (n=3). (D) Acetate, lactate, and succinate were not detected, as these organic acids are consumed by *R. palustris* Nx (LaSarre *et al.*, 2017). ****, statistical difference from corresponding Nx+WT value (P<0.0001), determined by two-way ANOVA with Sidak posttest. (A-E) Error bars, SD. (B, C, E) Statistical differences were determined using an unpaired, two-tailed t test; ns, non-significant.

Since all growth and metabolic trends we assayed were statistically similar between the Nx+WT and Nx+ΔFdhF cocultures, except for formate and H_2_ yields, we reasoned that the formate and H_2_ yields were unlikely to be influenced in any major way by factors other than the absence of *fdhF*. Thus, we estimated the contribution of *E. coli* to H_2_ production in coculture to be the difference in the H_2_ yield between the Nx+WT and Nx+ΔFdhF cocultures. From this difference, we estimated that *E. coli* produces 32±5% (SD) of the H_2_ in an Nx+WT coculture.

### Higher NH_4_^+^ excretion by *R. palustris* results in a larger *E. coli* contribution to H_2_ production

In the above cocultures with *R. palustris* Nx, the growth rates of the two species are coupled (LaSarre *et al.*, 2017). Consequently, *E. coli* grows at ~20% of the growth rate that would be possible if NH_4_^+^ were saturating, decreasing the rate of formate accumulation and acidification of the culture medium (LaSarre *et al.*, 2017). However, it is possible to uncouple the growth rates and allow *E. coli* to grow faster by growing *E. coli* with the hyper-cooperative *R. palustris* NxΔAmtB strain (LaSarre *et al.*, 2017). *R. palustris* NxΔAmtB lacks high-affinity AmtB transporters responsible for NH_4_^+^ import and thus excretes 3-times more NH_4_^+^ than does *R. palustris* Nx (LaSarre *et al.*, 2017). The higher level of NH_4_^+^ cross-feeding increases the *E. coli* growth rate in coculture and causes the rate of organic acid production by *E. coli* to exceed the rate of organic acid consumption by *R. palustris*. As a result, consumable organic acids accumulate along with formate and prematurely acidify the coculture and inhibit *R. palustris* growth unless the buffering capacity of the medium is raised. However, even without additional buffer, the coculture maintains reproducible trends through serial transfers (LaSarre *et al.*, 2017). The conversion of formate to H_2_ and CO2 by *E. coli* FHL requires anaerobic conditions and a pH below 7 and is influenced by the formate concentration (Rossmann *et al.*, 1991, Pinske & Sawers, 2016). Thus, we hypothesized that the faster growth of *E. coli* in coculture with *R. palustris* NxΔAmtB, and the associated acceleration of culture acidification and formate accumulation, might trigger earlier FHL activity and thereby increase *E. coli’s* contribution to H_2_ production.

We compared growth and metabolic trends in cocultures pairing *R. palustris* NxΔAmtB with either WT *E. coli* (NxΔAmtB+WT) or the ΔFdhF mutant (NxΔAmtB*+*ΔFdhF). Again, growth trends in cocultures with *R. palustris* NxΔAmtB were not significantly affected by the absence of *E. coli fdhF* (Fig. 4A-C). As expected, compared to cocultures with *R. palustris* Nx, cocultures with *R. palustris* NxΔAmtB reached stationary phase more quickly (Fig. 3A vs. Fig 4A) due to the increased growth rate of *E. coli* (LaSarre *et al.*, 2017). To take into account the shortened growth period of *R. palustris* NxΔAmtB-containing cocultures in our comparisons with *R. palustris* Nx-containing cocultures, we sampled *R. palustris* NxΔAmtB-containing cocultures at two time points: (i) 96 h, which roughly matches the time that *R. palustris* Nx-containing cocultures spent in stationary phase; and (ii) 167 h, which corresponds to the total time for coculture experiments with *R. palustris* Nx.

**Fig. 4.**
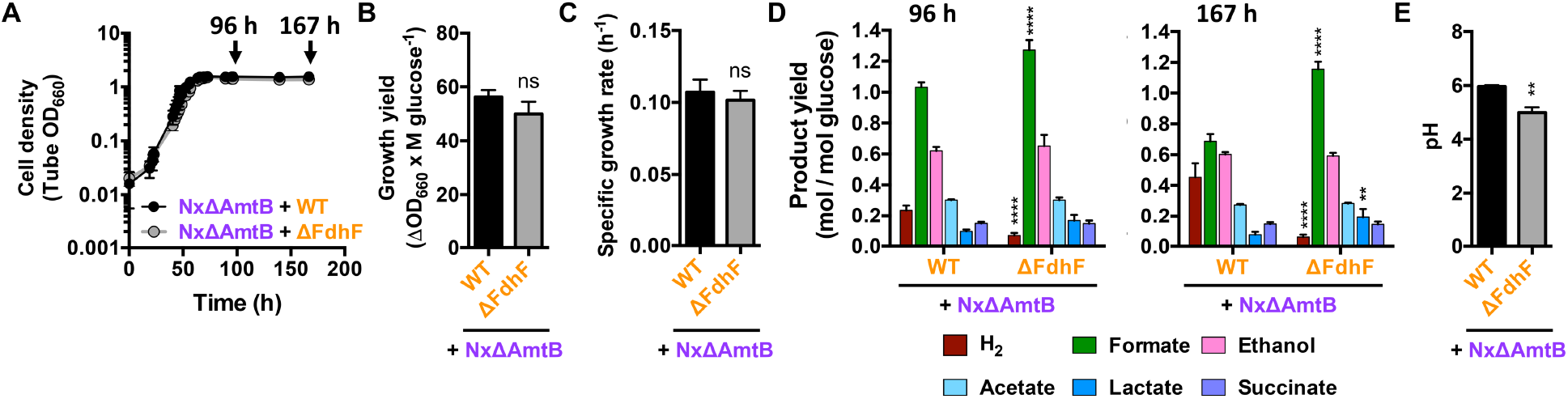
Growth and metabolic trends of cocultures pairing *R. palustris* NxΔAmtB with either WT *E. coli* or the ΔFdhF mutant. Coculture growth curves (A), growth yields (B), growth rates (C), product yields (D), and final pH (E) (n=3). (D) Asterisks indicate a statistical difference from the corresponding NxΔAmtB+WT value, with P<0.0001 (****) or P<0.01 (**), determined by two-way ANOVA with Sidak posttest. (A-E) Error bars, SD. (B, C, E) Statistical differences were determined using an unpaired, two-tailed t test; ns, non-significant; P<0.01 (**).

As observed previously (LaSarre *et al.*, 2017), consumable organic acids accumulated in cocultures with *R. palustris* NxΔAmtB (Fig. 4D). The average H_2_ yield of the NxΔAmtB+ΔFdhF cocultures across both time points was 0.10±0.01 mol/mole glucose (Fig. 4D), which is approximately one-third of that of the Nx+ΔFdhF cocultures (Fig. 3D). This H_2_ yield, which reflects the contribution by *R. palustris* NxΔAmtB, is likely low due to the inhibition of *R. palustris* growth and metabolism by acidification of the coculture before all consumable organic acids could be consumed (Fig. 4D and E), as observed previously (LaSarre *et al.*, 2017). In contrast, the NxΔAmtB+WT cocultures showed an increasing H_2_ yield between the two time points, eventually reaching levels comparable to those observed in Nx+WT cocultures (Fig. 4D). Taking the difference between the H_2_ yields of the NxΔAmtB+ΔFdhF and the NxΔAmtB+WT cocultures, we estimated that *E. coli* contributed 70±20 *%* and 86±26 *%* of the total H_2_ observed at 96 and 167 hours, respectively. Thus, unlike in Nx+WT cocultures, *E. coli* generated the majority of the H_2_ in NxΔAmtB+WT cocultures. This higher percent contribution by *E. coli* to H_2_ production was due in part to inhibition of *R. palustris,* but it was also due to *E. coli* having produced 2.4-times as much H_2_ per glucose in cocultures with *R. palustris* NxΔAmtB than in cocultures with *R. palustris* Nx (Fig. 5).

**Fig. 5.**
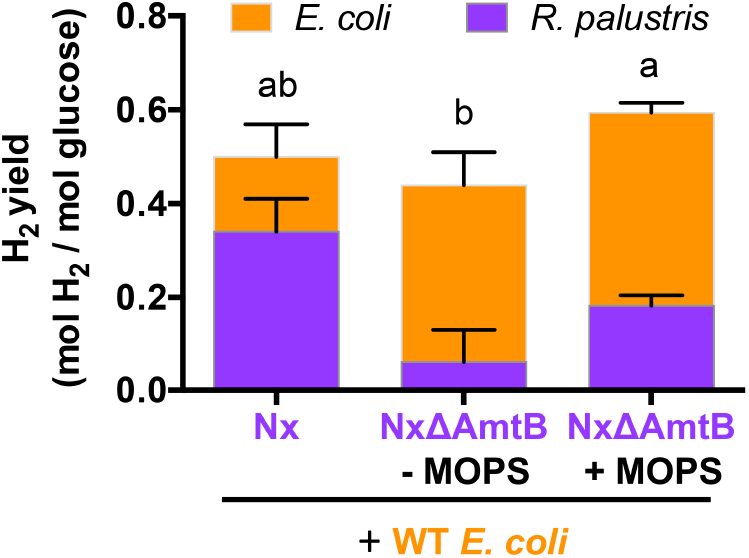
Species-level comparison of H_2_ yields in coculture. Yields were determined at the final time points shown in Figures 3, 4, and 6. The contribution of each species is estimated from the difference between cocultures with WT *E. coli,* which produces H_2_, and those with *E. coli* ΔFdhF, which does not produce H_2_. Different letters indicate a statistical difference between total H_2_ yields (P<0.05), determined using one-way ANOVA with Tukey’s multiple comparisons posttest.

The increase in H_2_ yield between time points in NxΔAmtB+WT cocultures corresponded with a decrease in the formate yield (Fig. 4D), indicating conversion of formate into H_2_ by WT *E. coli.* This formate removal also explains why the pH was higher in NxΔAmtB+WT cocultures compared to NxΔAmtB+ΔFdhF cocultures (Fig. 4E). The final pH of the NxΔAmtB+WT cocultures was also higher than that observed in a previous study on NxΔAmtB+WT cocultures (LaSarre *et al.*, 2017). Again, this difference is likely due to the removal of additional formate during the prolonged incubation; in the current study we sacrificed cultures to measure pH at 167 h (Fig. 4E), whereas previously we sacrificed cultures to measure pH at 96 h (LaSarre *et al.*, 2017). The extended incubation time and difference in the final pH between the two cocultures might also explain why the lactate yield was higher in the NxΔAmtB+ΔFdhF cocultures, because a low pH and fermentative conditions are known to stimulate lactate dehydrogenase activity in *E. coli* (Mat-Jan *et al.*, 1989, Jiang *et al.*, 2001).

The acidification of the medium in cocultures with *R. palustris* NxΔAmtB leaves some electron-containing organic acids unconsumed that *R. palustris* could otherwise convert to H_2_ via nitrogenase. To determine how much additional H_2_ could be made if *R. palustris* NxΔAmtB was not inhibited by the low pH, we repeated the experiments in medium supplemented with 100 mM MOPS, pH 7. This additional MOPS was not expected to inhibit *E. coli* FHL activity given that the pH of the medium without MOPS also has at a pH of 7, and both media acidify as *E. coli* grows fermentatively. Growth trends were similar between NxΔAmtB+WT and NxΔAmtB+ΔFdhF cocultures supplemented with MOPS (Fig. 6A-C). The presence of consumable organic acids at 94 h indicated that *E. coli* again grew rapidly and produced organic acids faster than *R. palustris* could consume them (Fig. 6D). However, the mildly acidic pH, only reaching 6.5 at 164 h (Fig. 6E), allowed *R. palustris* to eventually metabolize nearly all consumable organic acids (Fig. 6D). From the difference in H_2_ yields between MOPS-supplemented NxΔAmtB+WT and NxΔAmtB+ΔFdhF cocultures, we estimated that *E. coli* generated 63±10% of the H_2_ in cocultures at 94 h, similar to the *E. coli* contribution at 96 hours in Nx+WT cocultures (Fig. 4D). Thus, the additional MOPS buffer did not have a major inhibitory effect on FHL activity. By 164 h, the *E. coli* H_2_ contribution increased to 69±6% of the H_2_ produced, even though both species generated H_2_ during this time; for comparison, the H_2_ yield increased 1.4-fold between 94 and 164 h due to *R. palustris* nitrogenase activity alone in NxΔAmtB+ΔFdhF cocultures (Fig. 4D). The lower percentage of H_2_ contributed by *E. coli* in MOPS-supplemented NxΔAmtB+ΔFdhF cocultures compared to cocultures without MOPS was a result of increased H_2_ production by *R. palustris* NxΔAmtB, as the *E. coli* H_2_ yield was estimated to be similar in NxΔAmtB+ΔFdhF cocultures with and without MOPS (Fig. 5). As *R. palustris* NxΔAmtB was not the major H_2_ contributor even when allowed to fully consume the consumable organic acids, we conclude that the early exposure of *E. coli* to formate under FHL-activating conditions allows *E. coli* to make a greater contribution to H_2_ production than *R. palustris* in NxΔAmtB+WT cocultures. However, one reason that *R. palustris* NxΔAmtB did not make as much H_2_ in MOPS-supplemented cocultures compared to *R. palustris* Nx in coculture is because *R. palustris* NxΔAmtB shifted the *E. coli* fermentation balance towards ethanol, increasing the ethanol yield more than 2-fold above that observed in Nx+WT cocultures (Fig. 3D vs Fig. 4D and 6D). Because *R. palustris* does not consume ethanol in coculture, the high ethanol yield detracted from the electrons that *R. palustris* could otherwise have devoted to H_2_ production.

**Fig. 6.**
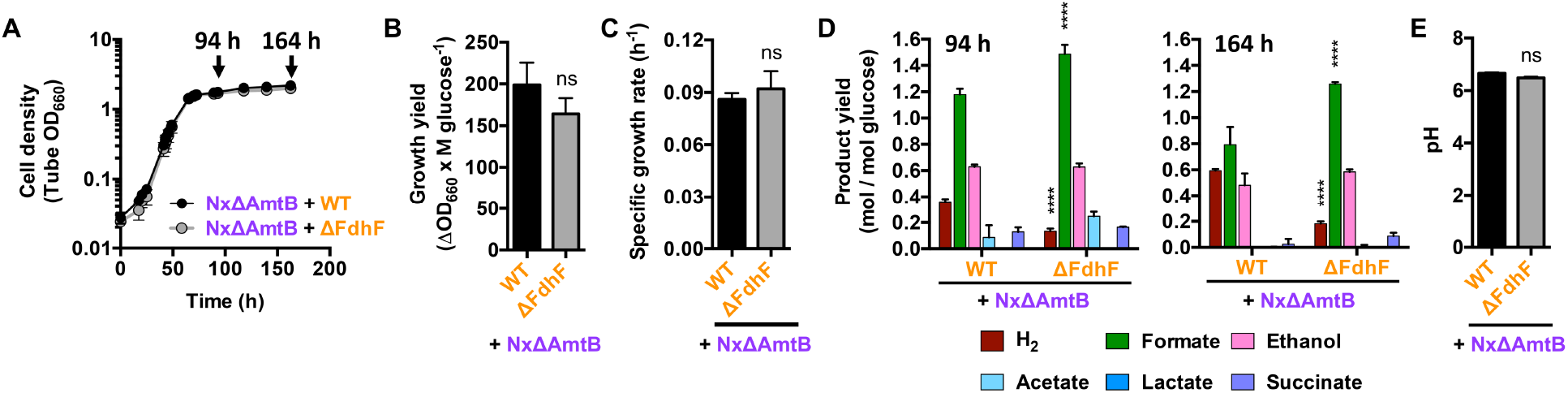
Growth and metabolic trends of cocultures pairing *R. palustris* NxΔAmtB with either WT *E. coli* or the ΔFdhF mutant in medium supplemented with 100 mM MOPS, pH 7. Coculture growth curves (A), growth yields (B), growth rates (C), product yields (D), and final pH (E) (n=3). (D) Asterisks indicate a statistical difference from the corresponding NxΔAmtB+WT value, with P<0.0001 (****), determined by two-way ANOVA with Sidak posttest. (A-E) Error bars, SD. (B, C, E) Statistical differences were determined using an unpaired, two-tailed t test; ns, non-significant.

## Discussion

Our results indicate that *E. coli* can make a substantial contribution to H_2_ production in cocultures with *R. palustris,* with the contribution ranging from 32-86% depending on the level of NH_4_^+^ excretion by the *R. palustris* partner and the length of time that *E. coli* is exposed to formate. Even in Nx+WT cocultures, wherein *E. coli* contributed the least (~32%) to H_2_ production (Fig. 3D), the contribution was still considerable in view of the fact that *E. coli* makes up only ~10% of the total population (LaSarre *et al.*, 2017, McCully *et al.*, 2017). This large contribution of *E. coli* to H_2_ on a ‘per cell’ basis reflects the difference in how electrons are managed in fermentation versus in photoheterotrophic growth. During fermentation, most of the electrons are disposed of in fermentation products, including H_2_, to satisfy electron balance. During photoheterotrophic growth by *R. palustris*, H_2_ production also contributes to electron balance, but most of the electrons are incorporated into new cell material (McKinlay & Harwood, 2010, McKinlay & Harwood, 2011). Thus, the relative biosynthetic efficiency of each species’ lifestyle plays a large role in determining its respective contributions to H_2_ production.

The H_2_ contribution by *E. coli* was greater in cocultures with *R. palustris* NxΔAmtB, in which the conditions required for *E. coli* FHL activity were established relatively early, thereby prolonging the period over which *E. coli* could convert formate to H_2_ (Fig. 4D and 6D). The greater *E. coli* contribution to H_2_ yields in NxΔAmtB+WT cocultures compared to that in Nx+WT cocultures could also be due in part to a larger *E. coli* population; *E. coli* makes up 30-50% of the total population in NxΔAmtB+WT cocultures, with absolute *E. coli* populations being ~2-fold larger in NxΔAmtB+WT cocultures compared to Nx+WT cocultures (LaSarre *et al*., 2017, McCully *et al*., 2017).

The results herein could contribute to the rational design of H_2_-producing communities. Much research has focused on the potential use of purple nonsulfur bacteria to convert fermented agricultural or municipal waste into H_2_. Coculture systems like ours can be viewed as a precursor for a consolidated process in which purple nonsulfur bacteria, like *R. palustris,* would be integrated with a fermentative community in situ. In our coculture, *E. coli* serves as a proxy for a fermentative community. While not all fermentative microbes generate H_2_ (Odom & Wall, 1983), our results show that fermentative bacteria could be major contributors to H_2_ production in communities with purple nonsulfur bacteria. Although the highest H_2_ yield observed in our study was 0.6 mol H_2_/mol glucose, or 5% of the theoretical maximum yield, it is possible that the yield would be higher if more time were allowed for *E. coli* to convert remaining formate into H_2_. Continuous removal of H_2_ from the headspace could also improve H_2_ production by relieving thermodynamic feedback on hydrogenase activity (Mandal *et al.*, 2006). It is also possible that the *R. palustris* contribution to H_2_ production could be increased by integrating *R. palustris* into a fermentative community in a manner where its access to nitrogen could be controlled, for example, using a latex biofilm (Gosse *et al.*, 2007, Gosse *et al.*, 2010, McKinlay & Harwood, 2010). We previously observed that nitrogen-starved *R. palustris* suspensions produced H_2_ at yields as high 66% of the theoretical maximum (McKinlay *et al.*, 2014). Similarly, in nitrogen-limited cocultures we observed an H_2_ yield of >4 mol H_2_/mol glucose, or 33% of the theoretical maximum yield (McCully *et al.*, 2017). Overall, our results illustrate how synthetic tractable communities can be used to inform on the design and application of microbial communities to benefit society.

